# A Fast and Low-Cost Approach for Binding Mode Validation of AI-Designed Therapeutics

**DOI:** 10.1101/2024.05.02.592265

**Authors:** Sara Zhang, Calvin Simmons, Mike Young, Jason Pan

## Abstract

High-resolution binding site mapping is important for in-depth activity assessment of new therapeutics including AI-designed antibodies. However, complex protein targets such as glycosylated antigens are challenging for many methods including crystallography. PD1 is a highly glycosylated antigen, and with the traditional HDX-MS method, only 51% sequence coverage could be obtained with multiple epitope residues undetected for Pembrolizumab. By implementing glyco-peptide detection, subzero temperature LC-MS and electron based MSMS fragmentation, the new HDX FineMapping methodology enabled 100% sequence coverage and complete epitope characterization for the Pembrolizumab-PD1 system, with amino acid level resolution. Furthermore, HDX FineMapping detects binding epitopes directly in solution, without any mutation or modification to either the antigen or the antibody. The amino acid level resolution combined with low cost, minimal sample consumption, fast turnaround time, and no need of mutant library or crystallization makes it a competitive methodology for binding mode validation of AI-designed therapeutics.

---

Glycosylation is an important post-translational modification (PTM) that occurs on most surface and secreted proteins in the cell,^1^ and it affects proteins folding, stability, and their biological activity. Due to the complex and dynamic nature of glycans, glycosylated protein antigens are challenging targets for many epitope mapping methods including X-ray crystallography, as the prerequisite crystal may not be obtained from the desired antibody-antigen complex. Programmed cell death protein 1 (PD1) has been an important drug target in recent years,^2,3^ and blockade of PD1 signaling pathway has enabled the successful development of anti-cancer monoclonal antibodies such as Pembrolizumab (Keytruda) and Nivolumab (Opdivo).^4,5^ The PD1 protein is highly glycosylated in human cells. With a calculated molecular weight of 16.8 kDa based on the amino acid sequence, it migrates as 33-38 kDa on reducing SDS-PAGE due to glycosylation. As shown in Figure 1, mass spectrometric measurement of intact full length PD1 molecules (expressed in HEK293) by LC-MS resulted in very complex mass peaks. Peaks a-g represent PD1 molecules with various number of glycan chains (up to 15 glycan units) on its glycosylation sites. It is also worth noting that the peak intensity of the non-glycosylated PD1 (non-glyco-PD1) is at the baseline, suggesting PD1 expressed in human cells are almost 100% glycosylated.

**Figure 1.**
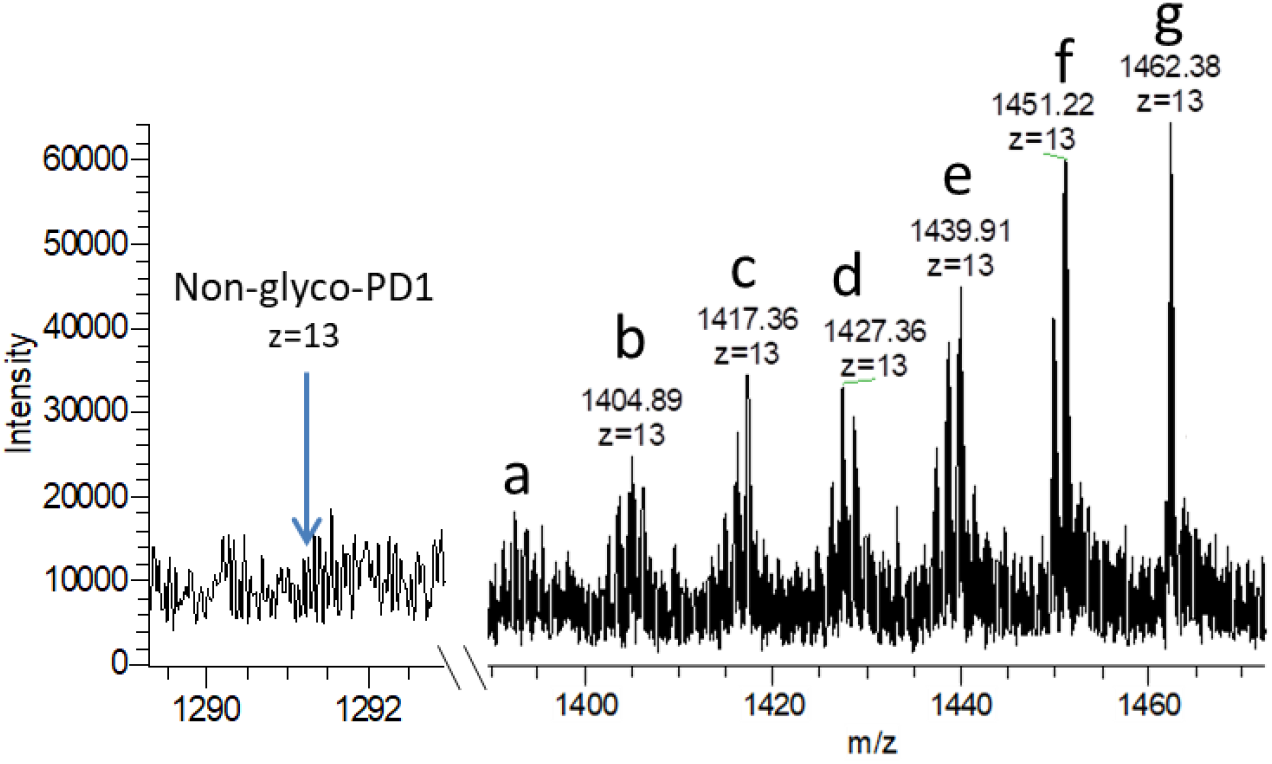
Mass spectrum of intact PD1 with various number of glycans.

Epitope mapping for glycosylated antigens are also challenging for methods other than crystallography. For example, with Pembrolizumab-PD1 system, the alanine substitution and binding affinity change based mutagenesis method (also called ‘alanine scanning’) revealed 10 residues out of 37 tested as the epitope, but only one residue (D85) is part of the actual epitope.^6^ Moreover, alanine substitution of 12 actual epitope residues had no binding loss at all, suggesting these epitope residues are undetectable (or ‘silent’) to the mutagenesis method. With the peptide array method, only the N-terminal epitope region was identified, which covered just five out of the actual 25 epitope residues of Pembrolizumab.^6^ With another chemical cross-linking method, also only five epitope residues could be detected. The authors also tested the traditional HDX-MS (hydrogen/deuterium exchange coupled with mass spectrometry) method for the Pembrolizumab-PD1 system,^6^ and it detected peptides covering 14 out of the actual 25 epitope residues, but no residue level information was available because the HDX-MS method used was only peptide-based and the lack of overlapping peptides. Moreover, the C-terminal epitope region was missed for unspecified reason.

PD1 is a relatively small protein, however, it contains five glycosylation sites as shown in Figure 2. In the traditional HDX-MS workflow, peptides derived from enzymatic digestion (usually pepsin) at acidic pH (2.5-3) must be identified first so that their H/D exchange behaviors can be followed in the next step.^7,8^ However, peptides with glycosylation are not taken into account in this step due to the complex nature of glycans. This is a big problem for glycosylated antigens like human PD1. Indeed, we got only 51% coverage of the PD1 sequence using the traditional HDX-MS method, leaving almost half of the protein undetected (Figure 2, top sequence). If this result is used for epitope mapping, any epitope information in that 49% region will be missed. Sometimes people claim they get the glycosylation site covered by detecting the nonglycosylated peptides in that region, even though the glycosylation percentage is >99%, thanks to the improved sensitivity of modern mass spectrometers. Unfortunately, however, this is not correct because the antibody developed against a glyco-antigen may not bind or bind differently to the nonglycosylated form. It has been reported that the binding of Nivolumab to human PD1 is dependent upon glycosylation of the antigen – it binds only to PD-1 expressed in a mammalian cell line but not to the nonglycosylated form expressed in E. coli (https://www.accessdata.fda.gov/drugsatfda_docs/nda/2014/125554Orig1s000PharmR.pdf). Therefore, examining the HDX behavior of peptides derived from the nonglycosylated form of a glyco-antigen may result in false epitope information.

**Figure 2.**
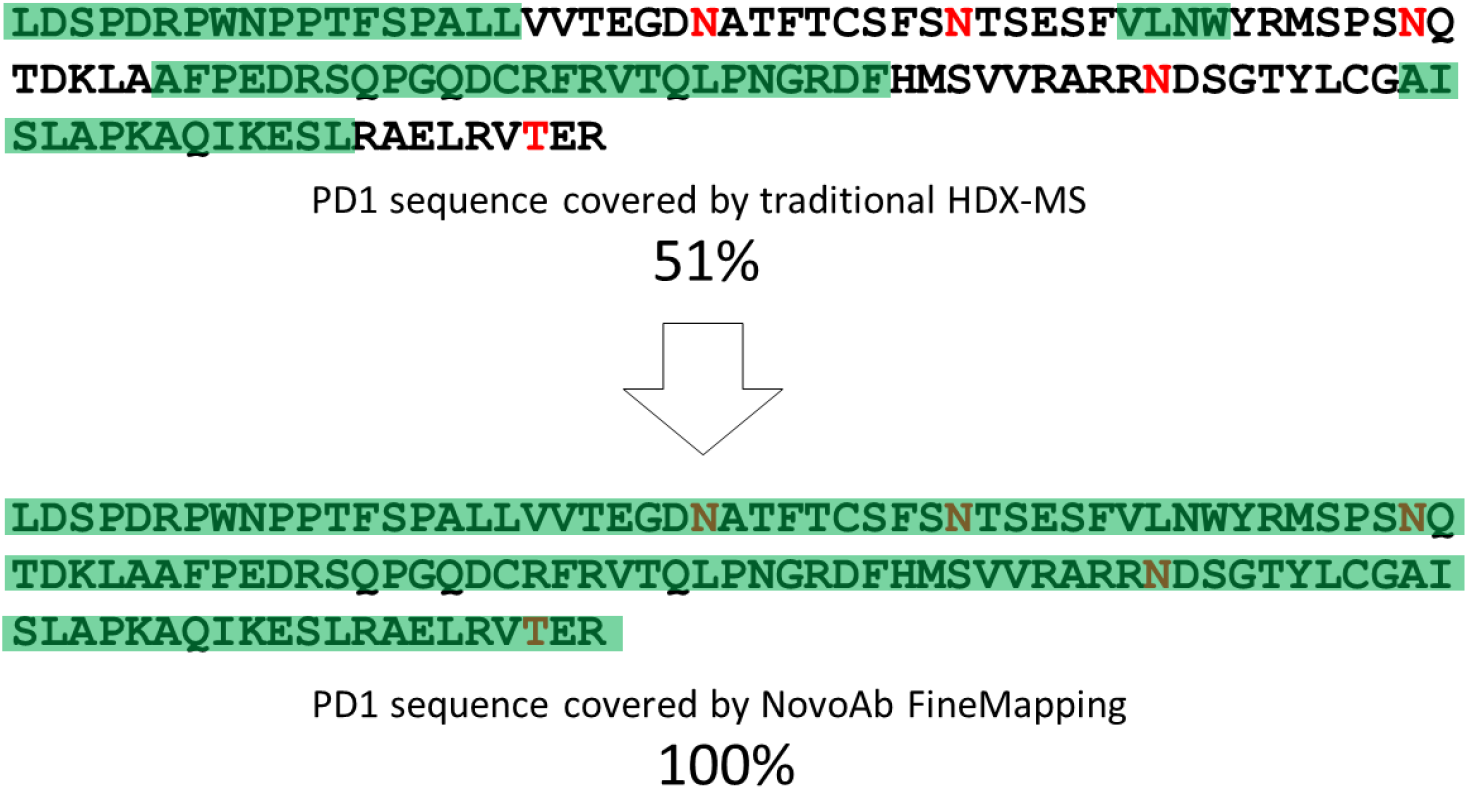
Sequence coverage by peptides obtained from traditional HDX-MS (51%, top) and from FineMapping (100%, bottom).

To overcome the limitations of traditional HDX-MS, here we developed a FineMapping methodology with advancements in three aspects: 1) direct identification of glyco-peptides using dedicated glycan search database followed by HDX examination; 2) LC-MS at subzero temperature (-20 °C); 3) an extra ETD-MSMS step on selected peptides. Subzero LC can reduce the D/H back exchange from >30% down to 2% as we reported previously,^9^ thus significantly enhance the epitope mapping sensitivity. On the other hand, the use of ETD fragmentation can enhance the resolution of HDX epitope mapping to single residue level. For the Pembrolizumab-PD1 system, full sequence coverage was obtained by this method, with all glycosylation sites covered (Figure 2, bottom sequence).

By measuring the deuterium uptake behavior of corresponding peptides with and without Pembrolizumab, HDX differential peptides were selected and subjected to ETD-MSMS under scrambling free conditions.^9-12^ An example of the ETD spectrum from a representative peptide is shown in Figure 3. ETD fragments covered the entire peptide and single residue resolution was achieved for both N-terminal c ions and C-terminal z ions. The deuterium content of each amino acid residue was calculated using the following equation, adapted from the analysis strategy we described previously,^9,10^

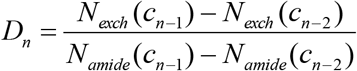

where *N*_*exch*_ is the normalized number of exchanged hydrogens for the corresponding fragment ion, while *N*_*amide*_ is the number of amide hydrogens included in the same fragment. Antibody binding induced protection is reflected on the H/D exchange reduction (ΔD) on the amino acid residues in the PD1-Pembrolizuma complex. Only those residues with a significant protection are assigned as epitope residues based on the criteria established previously. ^9,10,13^ Consequently, the epitope residues of Pembrolizumab are determined to be S60-Y68, Q75-K78, A81-G90, L128, and K131-A132 on PD1. It is worth noting that these residues covered almost all of the epitope residues determined by X-ray, where the nonglycosylated PD1 was used as the antigen for crystallization purposes (Figure 4).^14^ The only difference is seen in the C-terminal region, where FineMapping pinpointed three epitope residues while six were reported in the above literature. When we checked another publication on Pembrolizumab-PD1 complex with X-ray crystal structure at even higher resolution (2.15 Å vs 2.9 Å),^15^ there are indeed only three epitope residues (L128, K131 and A132) the C-terminal region, which are the same as determined by FineMapping. The three extra residues reported by the lower resolution X-ray article were found to be in no contact with Pembrolizumab.

**Figure 3.**
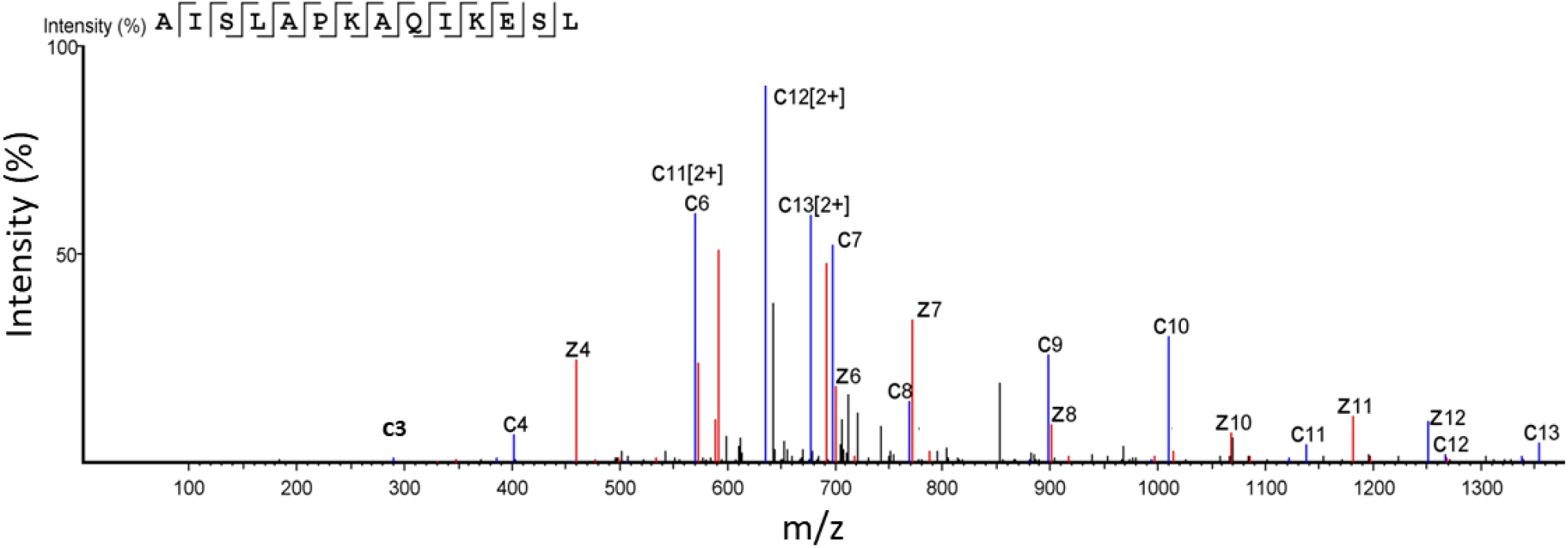
ETD-MSMS spectrum of a representative peptide from PD1.

**Figure 4.**
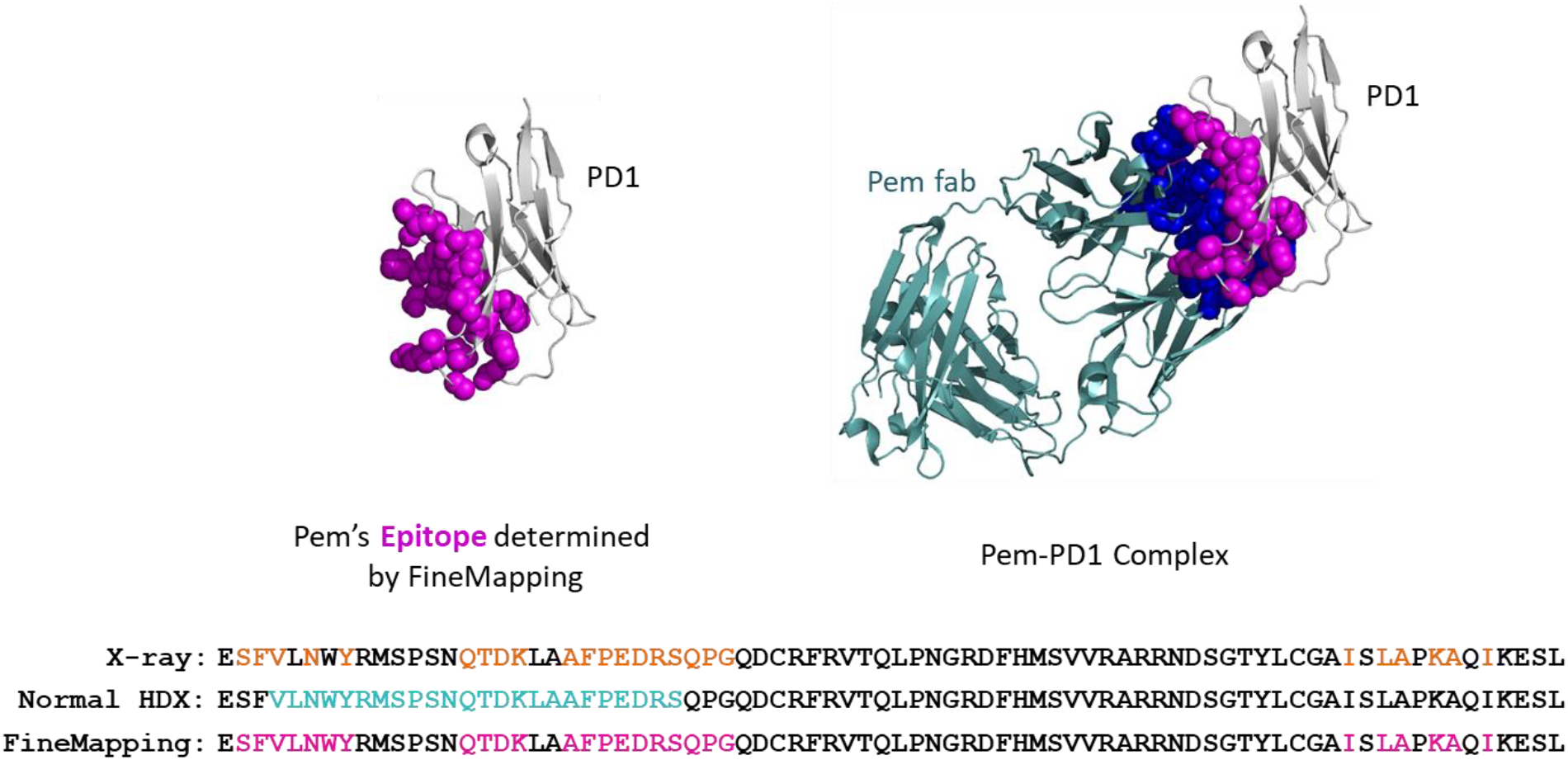
The epitope of Pembrolizumab on PD1 determined by NovoAb FineMapping (pink spheres). Sequence alignment at bottom: X-ray result is from reference 14, normal HDX result is from reference 6.

The location of FineMapping determined epitope residues on PD1 is shown in Figure 4. We also compared our results with literature data,^6^ where the traditional peptide based HDX-MS was used. The epitope thus defined was a long segment of the PD1 protein, without any residue level information (Figure 4, highlighted in cyan color). Moreover, 11 actual epitope residues of Pembrolizumab were missed probably due to lack of peptides from those glycosylated regions and a high degree of back exchange.^6^ In the FineMapping technology, we think the inclusion of glyco-peptides (to increase sequence coverage), running peptide LC separation at -20 °C (to reduce H/D back exchange), and an extra ETD-MSMS fragmentation step on epitope peptides (to get deuteration information on individual amino acid), are the main factors contributed to complete epitope characterization of Pembrolizumab with single residue level resolution. It is noteworthy that the detection of full epitope without any missed residues is a crucial step towards antibody characterization, not only for better understanding of how they interact with their target, but also for quality control and enhanced intellectual property (IP) protection. For example, the applicants of U.S. patent 9,115,188 successfully distinguished their mAb from the prior-art mAb by showing that the claimed mAb binds to a conformational epitope containing at least one amino acid not involved in the prior-art mAb’s epitope.

As the majority of proteins (about 50-70%) in the cell are glycosylated,^1^ the middle-down HDX-MS based FineMapping technology presented here should find its wide application on these protein targets. In fact, one can easily evaluate if their protein of interest is glycosylated or not by performing a quick search in the Uniprot database (https://www.uniprot.org, make sure to look at the results from Homo sapiens [human]). Also, FineMapping detects antibody epitopes directly in solution, without any mutation or modification to either the antigen or the antibody. The amino acid level resolution combined with low cost, minimal sample consumption, fast turnaround time, and no need of mutant library or crystallization makes it a competitive methodology for binding mode validation of AI-designed therapeutics.

